## Supplementary material for "eIF3 engages with 3’-UTR termini of highly translated mRNAs": Original gels

**Figure 1B**

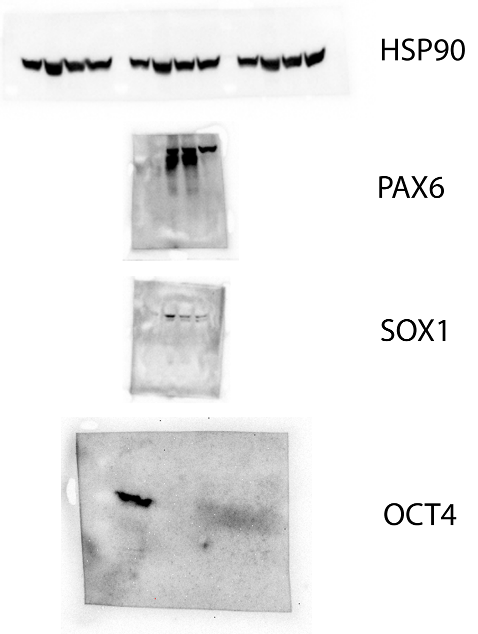


**Figure 1C**

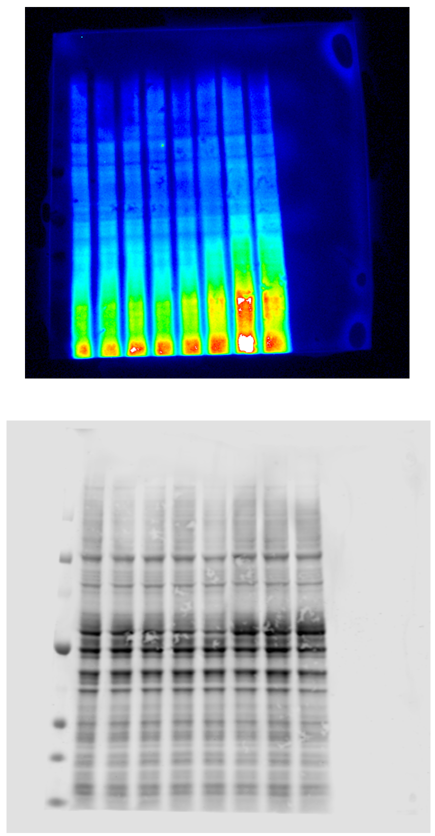

**Figure 1E**

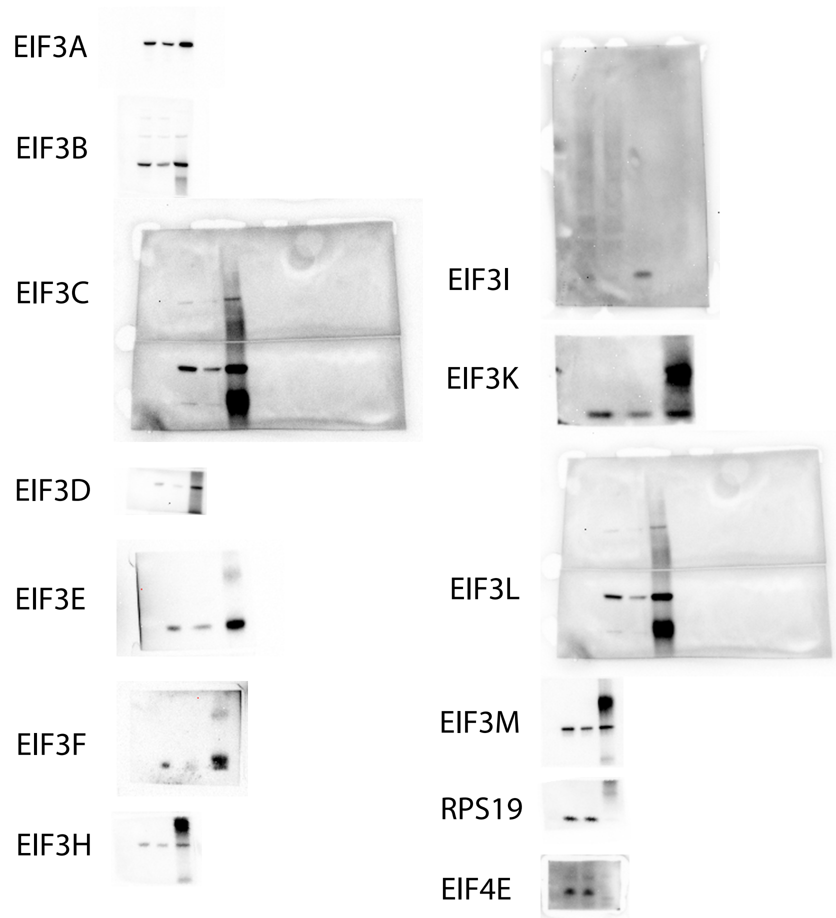


**Figure 1F**

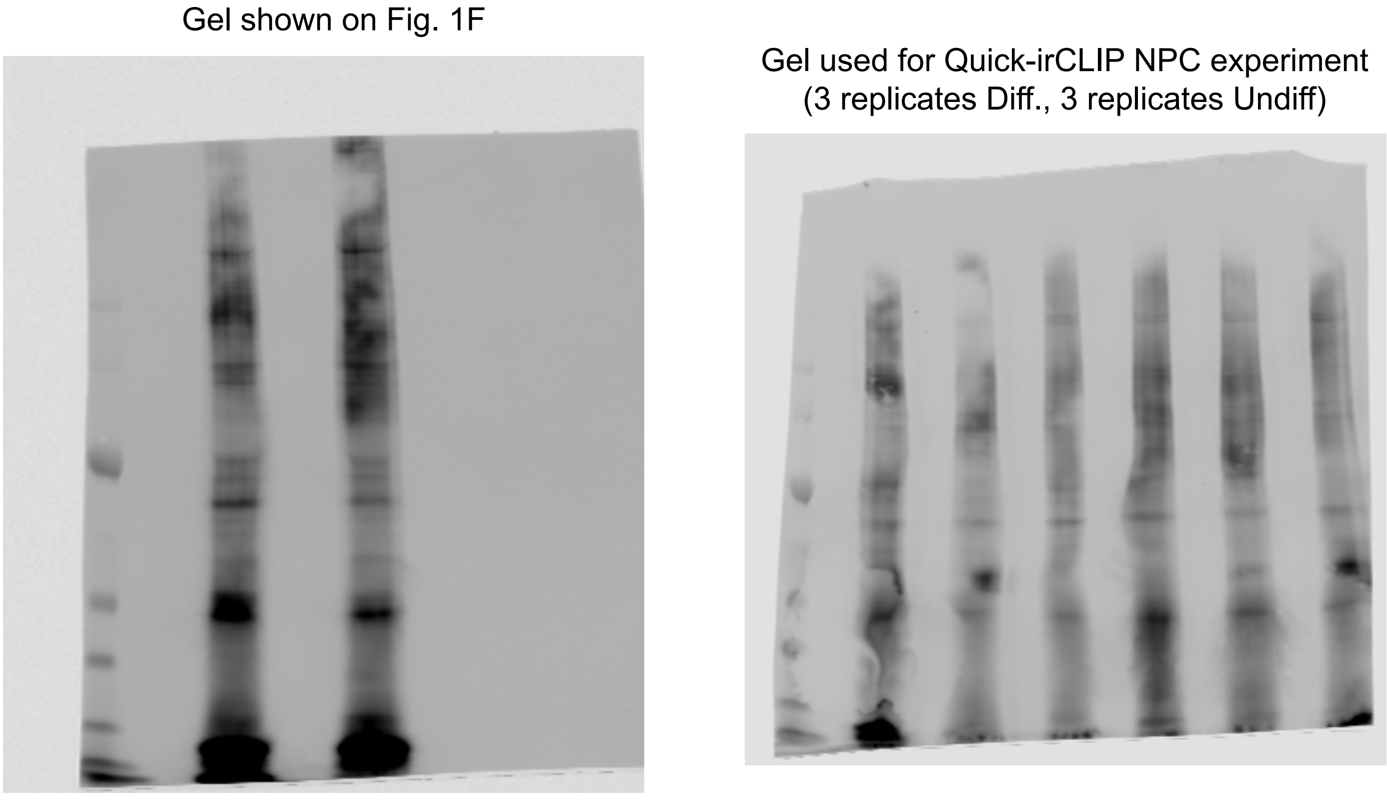


**Figure 3D**

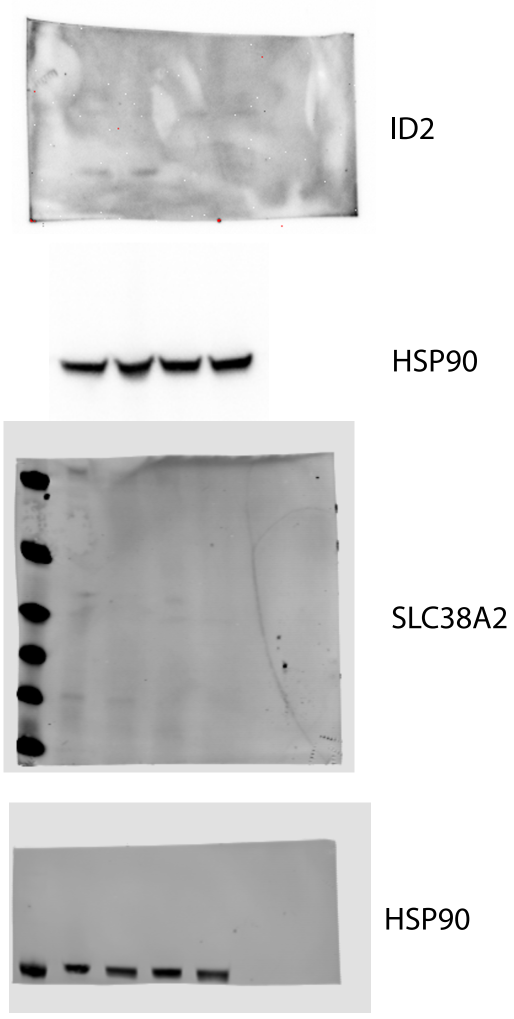


**Figure 3-figure supplement 3**

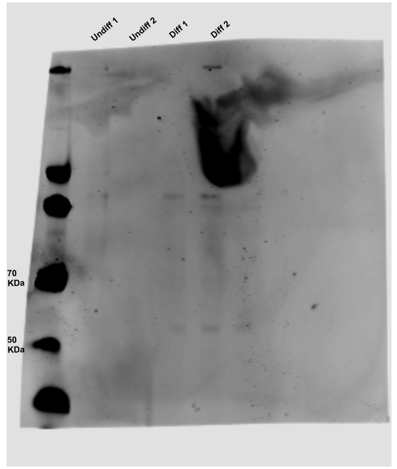

**Figures 4B and Figure 4-figure supplement 1**

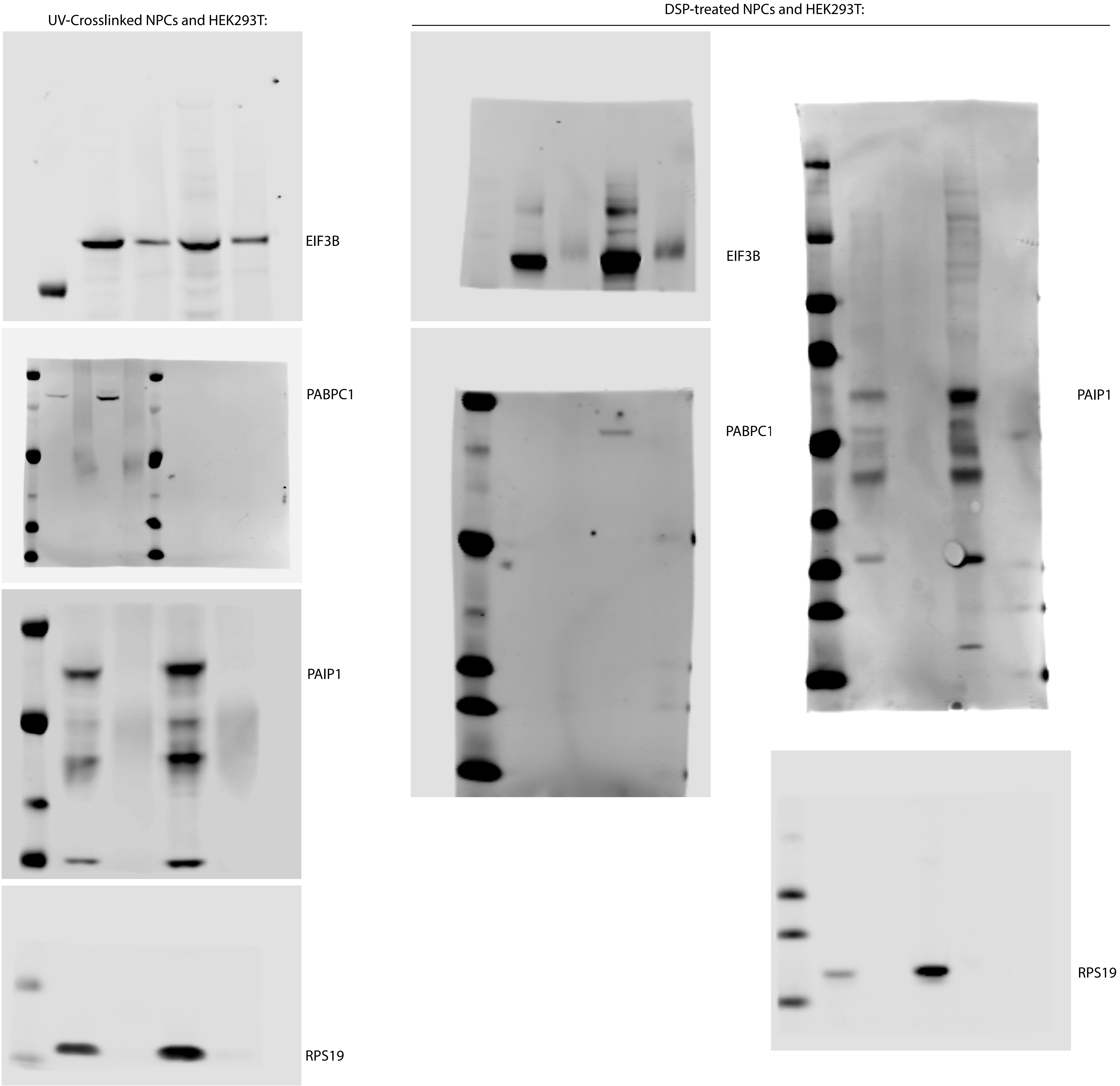
